# Functional characterization of SARS-CoV-2 vaccine elicited antibodies in immunologically naïve and pre-immune humans

**DOI:** 10.1101/2021.05.29.445137

**Authors:** David Forgacs, Hyesun Jang, Rodrigo B. Abreu, Hannah B. Hanley, Jasper L. Gattiker, Alexandria M. Jefferson, Ted M. Ross

## Abstract

As the COVID-19 pandemic continues, the authorization of vaccines for emergency use has been crucial in slowing down the rate of infection and transmission of the SARS-CoV-2 virus that causes COVID-19. In order to investigate the longitudinal serological responses to SARS-CoV-2 natural infection and vaccination, a large-scale, multi-year serosurveillance program entitled SPARTA (SARS SeroPrevalence and Respiratory Tract Assessment) was initiated at 4 locations in the U.S. The serological assay presented here measuring IgG binding to the SARS-CoV-2 receptor binding domain (RBD) detected antibodies elicited by SARS-CoV-2 infection or vaccination with a 95.5% sensitivity and a 95.9% specificity. We used this assay to screen more than 3100 participants and selected 20 previously infected pre-immune and 32 immunologically naïve participants to analyze their antibody binding to RBD and viral neutralization (VN) responses following vaccination with two doses of either the Pfizer-BioNTech BNT162b2 or the Moderna mRNA-1273 vaccine. Vaccination not only elicited a more robust immune reaction than natural infection, but the level of neutralizing and anti-RBD antibody binding after vaccination is also significantly higher in pre-immune participants compared to immunologically naïve participants (*p*<0.0033). Furthermore, the administration of the second vaccination did not further increase the neutralizing or binding antibody levels in pre-immune participants (*p*=0.69). However, ~46% of the immunologically naïve participants required both vaccinations to seroconvert.

## Introduction

In December 2019, individuals linked to an animal market in Wuhan, China presented with what was initially described as atypical pneumonia. Bronchoalveolar lavage fluid collected from hospitalized individuals contained a novel coronavirus detected by Illumina and Nanopore sequencing and electron microscopy^1^. This novel coronavirus, SARS-CoV-2, is a member of the *Betacoronavirus* genus in the *Coronaviridae* family^2^. Four main structural proteins make up the SARS-CoV-2 virion: nucleocapsid proteins surround the positive strand RNA genome, membrane proteins connect the membrane to the nucleocapsid, envelope proteins facilitate budding and detachment from the host cell, and spike proteins are involved in host receptor binding^3^. Similar to the closely related SARS virus that caused a 2003 outbreak, SARS-CoV-2 targets the ACE2 receptor located on the surface of the host cells^4–7^.

Since its initial identification, SARS-CoV-2 spread rapidly and resulted in a global pandemic. According to the World Health Organization, there is a cumulative burden of over 168 million confirmed cases and 3.5 million deaths as of May 28, 2021. In order to effectively combat the ongoing pandemic, a combination of targeted interventions and effective vaccines are required in conjunction with a deeper understanding of the elicited immune responses. The SARS-CoV-2 spike protein has been used as the primary antigen in several vaccines authorized for emergency use (EUA) by the U.S. Food and Drug Administration (FDA)^8–10^. The spike protein is highly immunogenic, and is able to elicit a wide array of serological and cellular responses^11,12^. Many neutralizing antibodies isolated from convalescent human sera specifically target the 223 amino acid receptor binding domain (RBD; amino acids 319-541) of the 1273 amino acid spike protein^13,14^. Therefore, RBD-directed antibodies are a suitable predictor of both serological binding and neutralizing potential^13,15–17^.

In December 2020, two mRNA-based vaccines received EUA in the U.S. Both the Pfizer-BioNTech BNT162b2 vaccine (2x 30μg doses 21 days apart) and the Moderna mRNA-1273 vaccine (2x 100μg doses 28 days apart) contain mRNA coding for the full-length spike protein^8,9^. These vaccines are delivered intramuscularly in a positively charged lipid nanoparticle to enhance host cell uptake^8,9^. Following endocytosis and endosomal escape, the mRNA is translated into protein by the host cells. Two proline mutations in the C-terminal S2 fusion machinery were inserted into the pre-fusion spike protein conformation to more closely mimic the intact virus^18–20^. When displayed by antigen-presenting cells (APCs), the spike protein stimulates a similar immune response to antigens presented over the course of a natural infection^21^. In human trials, both vaccines reported 95% efficacy in healthy adults in preventing symptomatic infection by SARS-CoV-2^8,9^. Full immunization is achieved 14 days following the second vaccination^9,22^. However, it is yet to be determined how quickly serological immunity wanes or what the exact nature of the memory response is upon re-exposure to the virus.

In order to explore these questions, our group initiated a large-scale longitudinal surveillance program entitled SPARTA to investigate the durability and effectiveness of immune responses to SARS-CoV-2 infection, re-infection, and vaccination. Blood was collected for the analysis of serological and cellular immune responses, and saliva was collected to test for the presence of SARS-CoV-2 viral RNA by nucleic acid amplification test (NAAT). Many SPARTA participants belong to high-risk groups prioritized for receiving early vaccination.

Here, we present validation for our serological assays used for SPARTA, as well as data tracking SARS-CoV-2-specific antibody levels in 32 immunologically naïve and 20 previously infected pre-immune participants going through vaccination by measuring both anti-RBD antibody binding via indirect ELISA, and viral neutralization (VN) using the USA-WA1/2020 SARS-CoV-2 strain. We also demonstrated how the effect of a single vaccination differs in immunologically naïve and pre-immune participants. While most researchers focus only on using ELISA or surrogate virus pseudo-neutralization assays as a substitute for VN with varying results^23–27^, we felt it was not only important to demonstrate the capacity of antibodies to bind to a SARS-CoV-2-derived protein, but also to assess *in vitro* protection against infectious SARS-CoV-2 virus by analyzing the potential of serum antibodies to limit virus-mediated cytopathic effects.

## Materials and Methods

### Ethics statement and the role of the funding source

The study procedures, informed consent, and data collection documents were reviewed and approved by the WIRB-Copernicus Group Institutional Review Board (WCG IRB #202029060) and the University of Georgia. The funding sources had no role in sample collection nor the decision to submit the paper for publication.

### SPARTA participant selection

Eligible volunteers between the ages of 18 and 90 years old (y.o.) were recruited and enrolled with written informed consent beginning March, 2020. Participants in the SPARTA program were enrolled at four locations: Athens, GA, Augusta, GA, Los Angeles, CA, and Memphis, TN in the U.S., and blood and saliva samples were collected monthly. Participants were predominantly people in high-risk groups, such as health-care workers, first responders, the elderly, and university employees and students receiving in-person tuition. Exclusion criteria included being younger than 18 years old, weighing less than 110 lbs, being pregnant, being cognitively impaired, or having anemia or a blood-borne infectious condition such as hepatitis C or HIV. As of May 28, 2021, 3124 participants were enrolled in the study and enrollment is ongoing. 69.9% of the participants identified as female and 29.2% identified as male. The mean age was 43.7 years (range 18-90). 76.4% of the participants self-identified as White/Caucasian, 11.7% as Black/African-American, 8.1% as Asian, and 3.8% as other or multiple.

Of those participants, 40 were chosen based on their vaccination and infection status as well as serum availability, and assorted into age and vaccine-matched groups based on the presence of SARS-CoV-2 infection prior to vaccination. At the pre-vaccination timepoint, 20 of them were categorized as immunologically naïve to SARS-CoV-2, and reported no specific COVID-19 symptoms, never had a positive NAAT result, and tested negative for RBD-specific IgG antibodies with concentrations below our experimentally determined threshold of 1.139μg/mL [see under *ROC analysis* heading] (Table S1A). The other 20 participants were immunologically pre-immune due to a previous SARS-CoV-2 infection prior to vaccination and reported COVID-19 symptoms, positive NAAT, and/or antibody concentrations above the threshold (Table S1B).

Serum samples were collected from 8 of the 20 pre-immune participants between the two vaccinations. In contrast, mid-vaccine serum was collected from only one of the 20 matched immunologically naïve participants. Therefore, an additional 12 immunologically naïve participants were selected who had a mid-vaccine timepoint, in order to statistically compare how a single vaccine dose affects antibody levels in immunologically naïve and pre-immune individuals (Table S1C). The mean age across all 52 participants was 45 y.o. (SD = 15.7, range = 24-72), with 57.7% female (n=30) and 42.3% male (n=22). 79% of the participants received the Pfizer-BioNTech BNT162b2 vaccine (n=41), while 21% received the Moderna mRNA-1273 vaccine (n=11) between December 18, 2020 and February 18, 2021.

### Blood collection and processing

BD Vacutainer serum separation venous blood collection tubes (BD, Franklin Lakes, NJ, USA) containing whole blood were centrifuged at 2500 rpm for 10 min. After centrifugation, the supernatant serum layer was isolated and heat inactivated in a 56°C water bath for 45 min to disable any infectious SARS-CoV-2 virus^28^. Serum was thereafter stored at −80°C.

### Enzyme-linked immunosorbent assay (ELISA)

Immulon^®^ 4HBX plates (Thermo Fisher Scientific, Waltham, MA, USA) were coated with 100 ng/well of recombinant SARS-CoV-2 RBD protein in PBS overnight at 4°C in a humidified chamber. Plates were blocked with blocking buffer made with 2% bovine serum albumin (BSA) Fraction V (Thermo Fisher Scientific, Waltham, MA, USA) and 1% gelatin from bovine skin (Sigma-Aldrich, St. Louis, MO, USA) in PBS/0.05% Tween20 (Thermo Fisher Scientific, Waltham, MA, USA) at 37°C for 90 min. Serum samples from the participants were initially diluted 1:50 and then further serially diluted 1:3 in blocking buffer to generate a 4-point binding curve (1:50, 1:150, 1:450, 1:1350), and subsequently incubated overnight at 4 °C in a humidified chamber. Plates were washed 5 times with PBS/0.05% Tween20 and IgG antibodies were detected using horseradish peroxidase (HRP)-conjugated goat anti-human IgG detection antibody (Southern Biotech, Birmingham, AL, USA) at a 1:4,000 dilution and incubated for 90 min at 37°C. Plates were then washed 5 times with PBS/0.05% Tween20 prior to development with 100μL of 0.1% 2,2’-azino-bis(3-ethylbenzothiazoline-6-sulphonic acid) (ABTS, Bioworld, Dublin, OH, USA) solution with 0.05% H2O2 for 18 minutes at 37°C. The reaction was terminated with 50μL of 1% (w/v) SDS (VWR International, Radnor, PA, USA). Colorimetric absorbance was measured at 414nm using a PowerWaveXS plate reader (Biotek, Winooski, VT, USA). All samples and controls were run in duplicate and the mean of the two blank-adjusted optical density (OD) values were used in downstream analyses. IgG equivalent concentrations were calculated based on a 7-point standard curve generated by a human IgG reference protein (Athens Research and Technology, Athens, GA, USA), and verified on each plate using human sera with known concentrations.

### ROC analysis and threshold determination using a validation cohort

In order to distinguish between antibody positive and negative participants, ROC (receiver operating characteristic) analysis was performed on a validation cohort comprised of 22 NAAT-confirmed positive and 49 pre-pandemic human sera. Serum from NAAT-confirmed positive participants were obtained from samples intended to be discarded from a central Georgia hospital in May and June 2020. They were all between the ages of 22 and 60 (mean = 43.5 y.o.) with a nearly even female (n=11) to male ratio (n=10) with one participant whose gender was unknown (Table S2). All participants tested positive for the presence of SARS-CoV-2 viral RNA collected via nasopharyngeal swab by NAAT 9-74 days prior to the blood draw. In addition, the presence of any COVID-19 symptoms the participants had experienced at the time of the NAAT were recorded (Table S2). These symptoms included, but were not limited to fever, cough, chills, loss of taste or smell, and shortness of breath. The pre-pandemic sera were collected between 2013 and 2018. The threshold between positives and negatives was chosen at the antibody concentration corresponding to the most similar sensitivity and specificity values, in order to balance false positive and false negative rates. Linear regression and correlation analyses were applied to test the relationship between anti-RBD IgG antibody concentrations (ELISA) and VN endpoint titers. All analyses were performed using GraphPad Prism 9.1.1.

### Viral neutralization assay

All research activities using infectious SARS-CoV-2 virus occurred in a Biosafety Level 3 (BSL-3) laboratory in the Animal Health Research Center at the University of Georgia (Athens, GA, USA). The USA-WA1/2020 SARS-CoV-2 strain was propagated as previously described^29^. 50μl of two-fold serially diluted serum (1:5–1:640 or 1:50–1:6400 depending on the magnitude of antibody binding concentration and vaccination status) was incubated with the SARS-CoV-2 virus (100 TCID50/50μl) for 1 h at 37°C. The serum-virus mixture was then transferred to Vero E6 cells in 96-well cell culture plates. The plates were observed for 3 days for cytopathic effects (CPEs), such as the aggregation and detachment of cells. The VN endpoint titer was determined as the reciprocal of the highest dilution that completely inhibited CPE formation.

### Statistical comparison of participant groups

Statistical difference between pre-, mid-, and post-vaccination timepoints for both the immunologically naïve and the infected cohorts were determined using a paired *t*-test, while the difference between the post-vaccination timepoints of immunologically naïve and infected participants was determined using an unpaired *t*-test. The pre-vaccination timepoints occurred at least 1 day prior to receiving the first dose of an mRNA vaccine (mean = 32.6 days, range = 1– 99 days), mid-vaccination timepoints occurred between the administration of the two vaccinations (mean = 13.5 days, range = 4-26 days), while post-vaccination timepoints occurred a minimum of 14 days after the administration of the second dose of the same mRNA vaccine (mean = 28.8 days, range = 4–51 days). Statistical analyses were performed using GraphPad 9.1.0, and statistical significance was represented by * *p*<0.05, ** *p*<0.01, *** *p*<0.001, and **** *p*<0.0001.

## Results

In order to establish the threshold value between participants with positive and negative RBD-binding IgG antibody levels, a ROC analysis based on serum from the validation cohort of 49 pre-pandemic negative participants and 22 confirmed NAAT positive participants was performed. The threshold between negative and positive anti-RBD IgG antibody concentrations was set at 1.139μg/mL, yielding a sensitivity of 95.5% and a specificity of 95.9% (Fig. 1A-B).

**Fig. 1:**
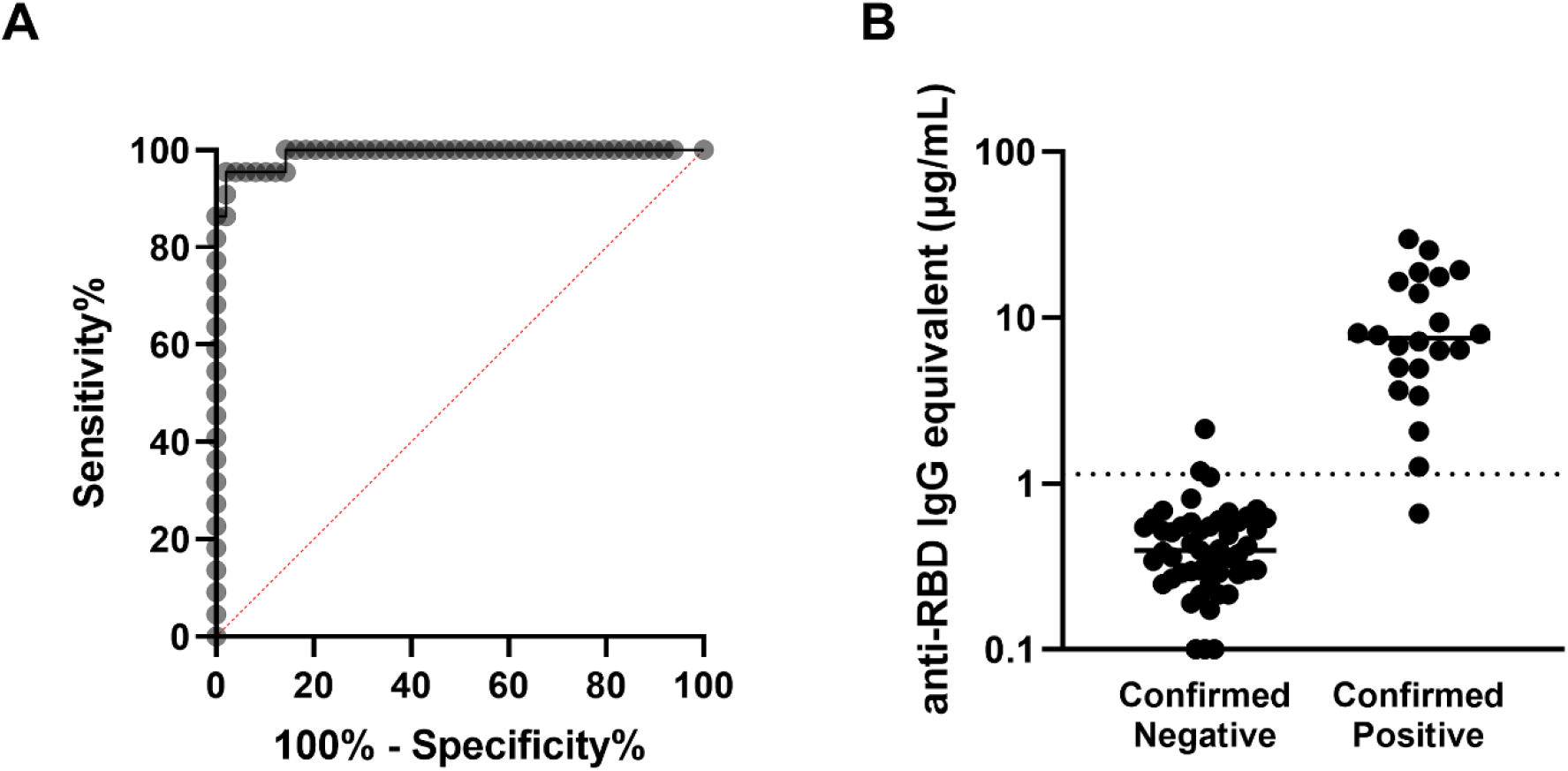
Determination of the antibody concentration threshold based on the validation cohort. (A) ROC analysis based on the 22 confirmed positive and 49 confirmed negative sera comprising the validation cohort. The area under the curve is 0.9917 (**** *p*<0.0001), and the sensitivity and specificity of this assay were determined to be 95.5% and 95.9% respectively, with the threshold between negative and positive set at 1.139μg/mL. (B) Anti-RBD antibody concentrations are shown for each participant in the validation cohort, sorted into negatives and positives based on prior confirmed COVID-19 status. The threshold is marked with the horizontal dotted line.

Further investigation into this validation cohort demonstrated that VN was absent in all participants with anti-RBD IgG antibody concentrations below the threshold, while all participants above the threshold showed some level of VN (Fig. 2A), and there was a strong correlation between ELISA binding and VN (*r*=0.9125, *p*<0.0001). Similarly, a strong positive relationship was observed between the antibody binding and neutralization levels for all timepoints of the 53 SPARTA participants from this study (*r*=0.9359, *p*<0.0001) (Fig. 2B).

**Fig. 2:**
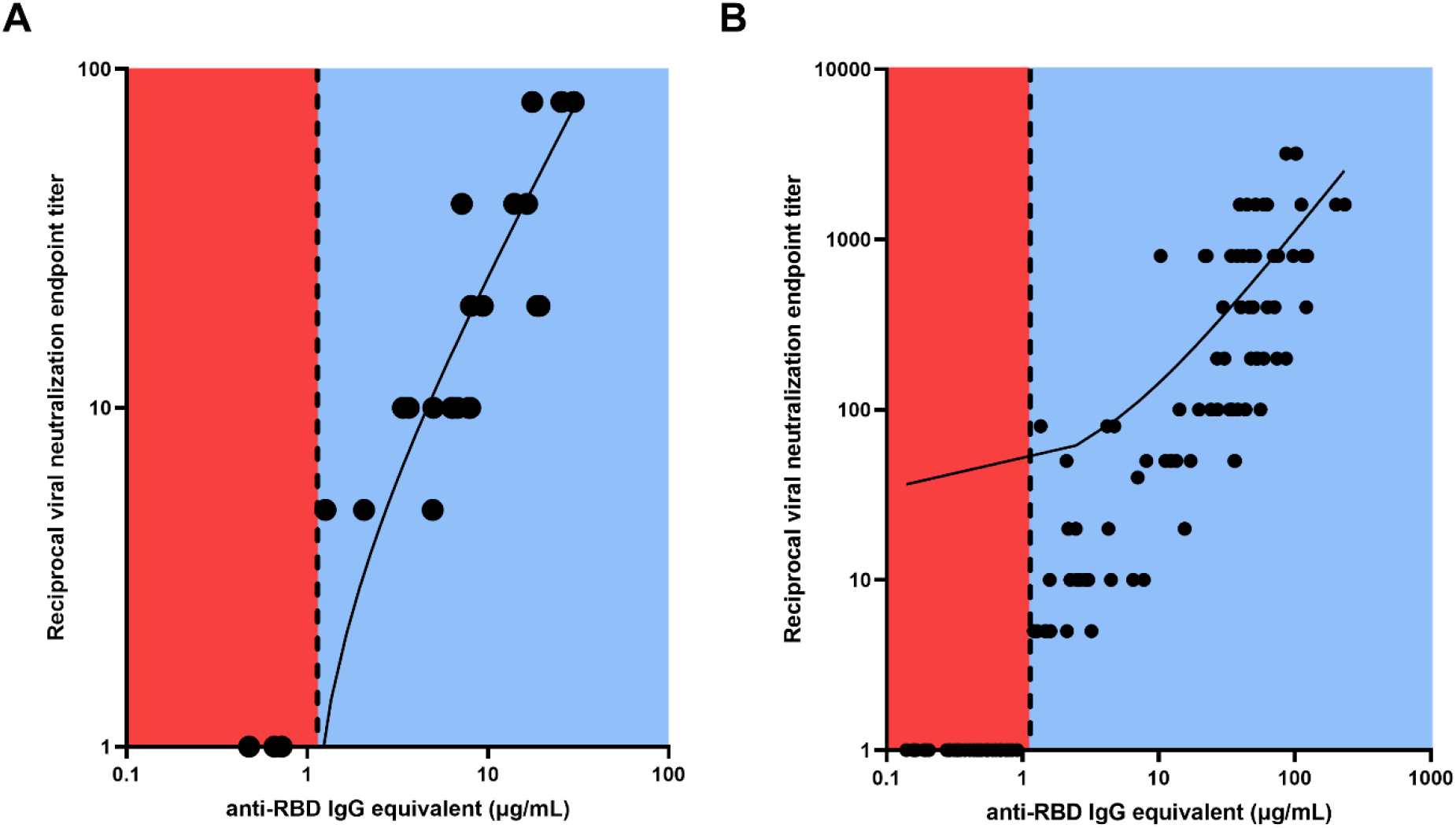
Linear regression between RBD-binding IgG antibody concentration and reciprocal VN endpoint titer. There was a strong positive relationship between antibody binding and VN titer for (A) the validation cohort (*r*=0.9125, **** *p*<0.0001), and (B) the SPARTA cohort (*r*=0.9359, **** *p*<0.0001). The vertical dotted line represents the threshold between negative and positive antibody concentrations, determined by the ROC analysis. No participants below the threshold demonstrated any neutralizing activity, while all participants above the threshold showed some level of VN. In order to represent the values on a logarithmic scale, lack of neutralization was reported as 1.

Using the experimentally determined threshold of 1.139μg/mL, 40 age-matched and vaccine-matched participants were selected, 20 of whom were immunologically naïve to SARS-CoV-2 prior to receiving a complete mRNA vaccine regimen, while the other 20 were pre-immune with signs of prior SARS-CoV-2 infection before vaccination. None of the immunologically naïve participants had any documented history of COVID-19, while 16 out of the 20 participants in the pre-immune group have reported COVID-19 symptoms or a positive NAAT. The remaining 4 were asymptomatic but had elevated RBD IgG antibody titers (Table S1A-B).

VN of the pre-vaccination and post-vaccination serum samples collected from these 40 participants demonstrated that vaccinations significantly increased the VN titer of both immunologically naïve and pre-immune participants (*p*<0.0001) (Fig. 3A). Interestingly, after vaccination, the neutralizing antibody titer of pre-immune individuals was significantly higher than the neutralizing antibody titer of immunologically naïve individuals (*p*<0.0001) (Fig. 3A). The post-vaccination neutralization titers for immunologically naïve participants ranged from 1:5–1:400, while the post-vaccination neutralization titers in pre-immune participants ranged from 1:400– 1:3200. The highest tested VN titer was 1:6400, but all participants demonstrated some level of CPE at this dilution point.

**Fig. 3:**
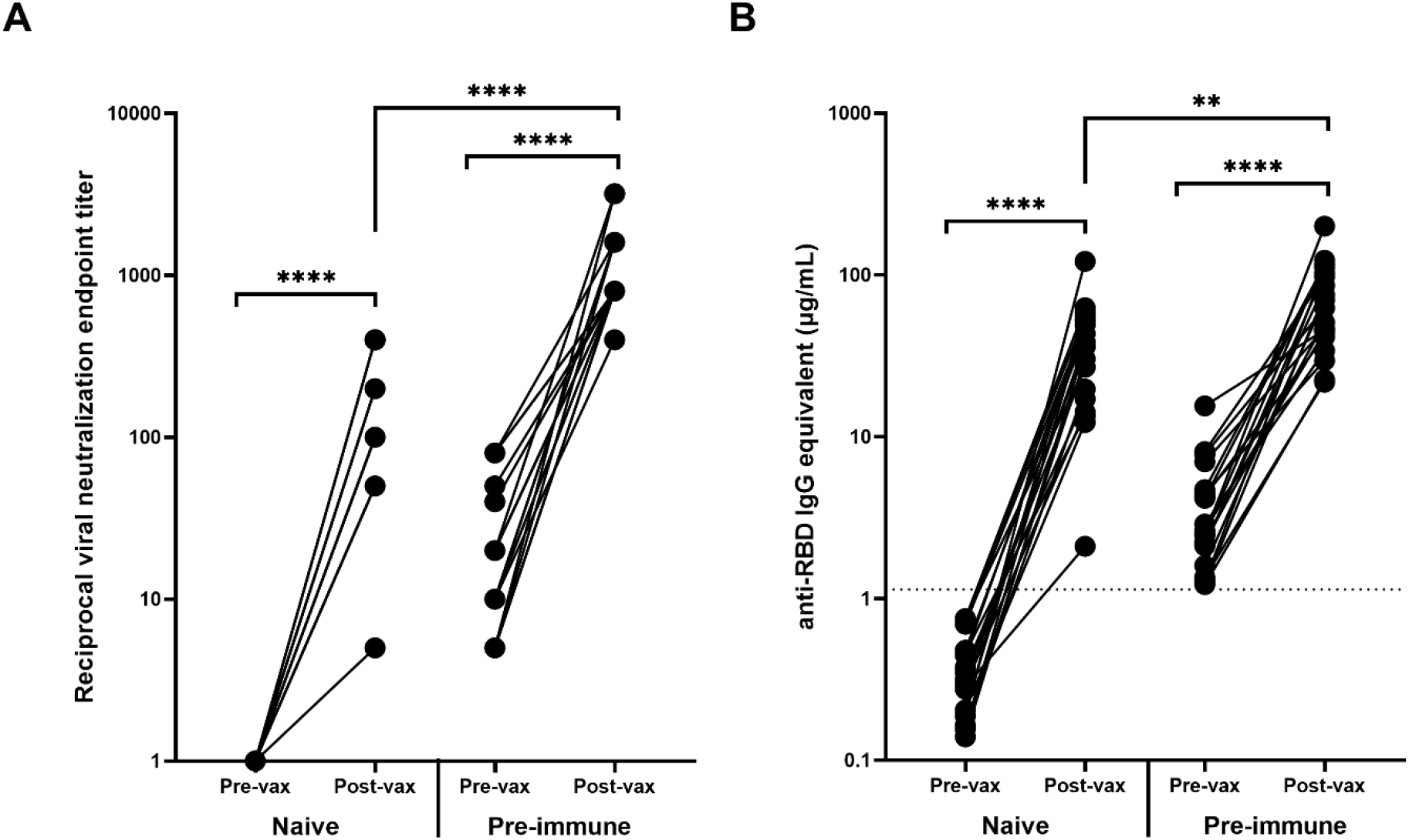
Antibody response before and after vaccination. Vaccination significantly increased the (A) VN and (B) RBD-binding antibody levels of all participants regardless of pre-vaccination naïve (n=20) or pre-immune (n=20) status (**** *p*<0.0001). The post-vaccination VN and anti-RBD IgG levels for participants who were pre-immune prior to vaccination were significantly higher (**** *p*<0.0001 and ** *p*=0.0033 respectively). Post-vaccination timepoints took place a minimum of 14 days after the administration of the second mRNA vaccination. In order to represent the values on a logarithmic scale, lack of neutralization was reported as 1.

Similarly, there were significant increases in RBD-binding antibody concentrations due to vaccination in both the immunologically naïve and pre-immune groups (*p*<0.0001) (Fig. 3B). The difference in post-vaccination antibody concentrations between immunologically naïve and pre-immune participants was less pronounced than demonstrated by VN titers, but antibody concentrations were still significantly higher in pre-immune participants (*p*=0.0033) (Fig. 3B).

Eight of the 20 participants in the age-matched pre-immune cohort had serum collected between the first and the second vaccinations (Table S1B). These timepoints ranged between 4 and 16 days after the first vaccination with a mean of 10.8 days (SD = 4.6). These previously infected participants had a significant increase in both VN titers (*p*=0.002) and RBD-binding IgG antibody concentrations (*p*=0.03) following the first vaccination, but there was no further significant change in RBD-binding or neutralizing antibody titers following the second vaccination (*p*=0.69) (Fig. 4A-B).

**Fig. 4:**
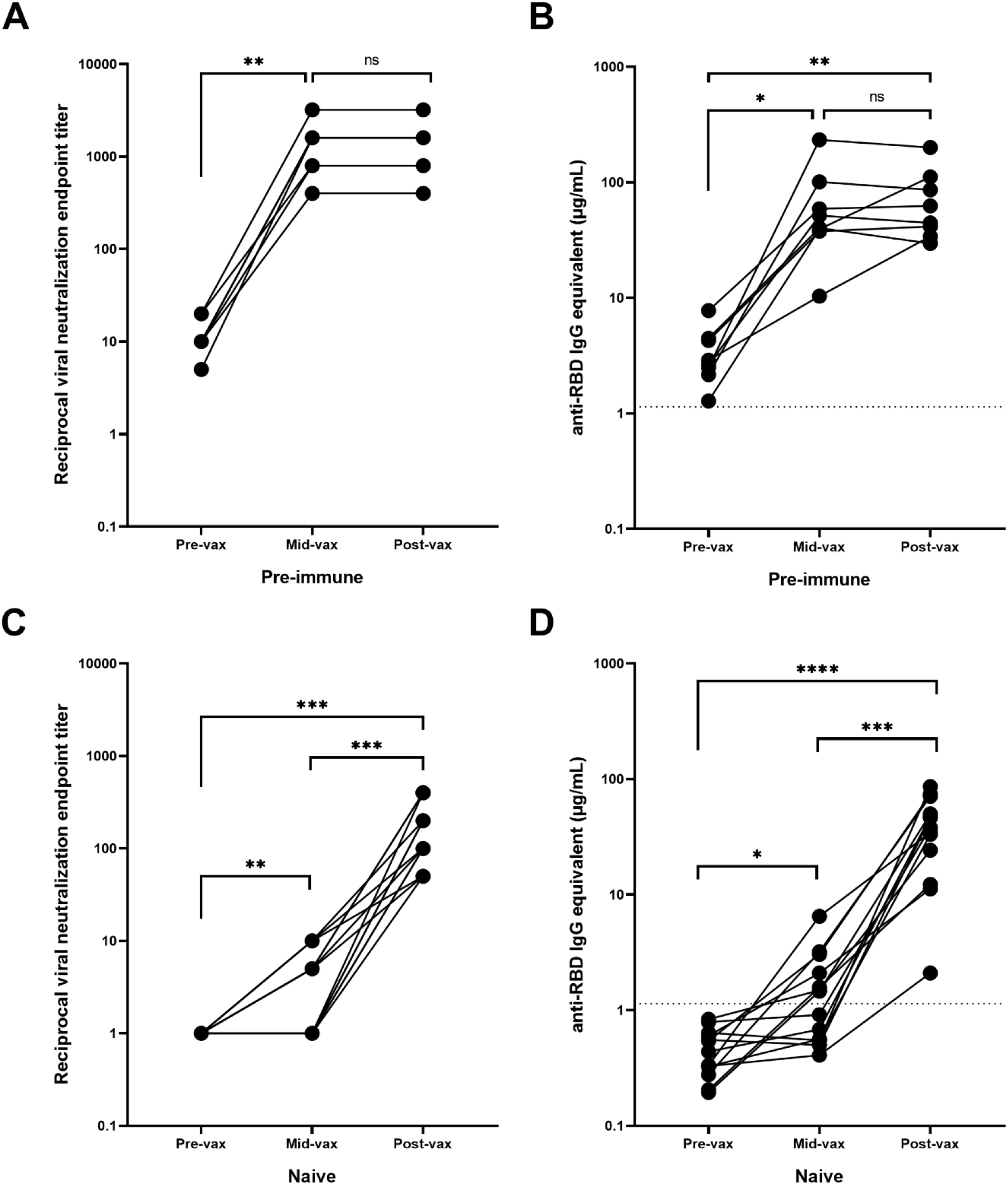
Antibody response during vaccination. Pre-immune participants (n=8) with serum obtained between the two mRNA vaccinations (mid-vax) have demonstrated a significant increase in (A) neutralizing (** *p*=0.002) and (B) anti-RBD antibody levels (* *p*=0.03) before the second vaccination was administered, and there was no significant change in antibody titer in response to the second vaccination (*p*=0.69). Immunologically naïve participants (n=13) with serum obtained between the two mRNA vaccinations have demonstrated a significant increase in (C) neutralizing (*** *p*=0.0005), and (D) binding antibody levels (**** *p*<0.0001). Six of them showed no neutralizing activity before the second vaccination was administered. While the other 7 showed a significant increase (** *p*=0.0074) and exhibited a low level of neutralization even after the first vaccination, the second vaccination was still crucial in boosting antibody levels (*** *p*=0.0006). In order to represent the values on a logarithmic scale, lack of neutralization was reported as 1.

Only one of the 20 naïve participants from the age-matched cohort had available samples collected between the two vaccinations (Table S1A). Therefore, an additional 12 participants were selected that had samples collected 9–26 days after the first vaccination (mean = 15.8 days, SD = 5.3 days), but prior to the second vaccination. Overall, there was a significant increase in neutralizing and RBD-binding antibody titers following both the first (*p*=0.0074 and *p*=0.02 respectively) and second vaccinations (*p*=0.0006 and *p*=0.0001 respectively) (Fig. 4C-D). However, 46% of these participants (n=6) had no detectable neutralizing activity or anti-RBD IgG antibodies after the administration of the first vaccination (mean = 12 days after the first vaccination, SD = 3.7 days, range = 9–18 days). These participants only seroconverted after receiving the second vaccination. The remaining participants (54%, n=7) exhibited a low level of positive neutralizing and binding antibody levels following the first vaccination (mean = 19 days after the first day, SD = 4.3 days, range = 12–26 days). There was a significant rise in neutralizing and anti-RBD IgG antibody levels following the second vaccination (*p*=0.023 and *p*=0.009 respectively).

## Discussion

Accurately tracking serological responses in people infected with or vaccinated against SARS-CoV-2 is critical in assessing the effectiveness of the induced immunity, which can subsequently inform public health decisions. Following SARS-CoV-2 infection, there is an expansion of B and T cells directed at various antigens in the virus, especially the receptor binding domain of the highly immunogenic spike protein^13,15–17^. Currently, COVID-19 vaccines authorized in the U.S. include the spike protein as the vaccine antigen that elicits protective immunity^8,9^. In this study, we examined the serological responses elicited by immunologically naïve or previously infected pre-immune individuals who were subsequently vaccinated with mRNA-based COVID-19 vaccines.

RBD was chosen over the full-length spike protein as the antigenic target for binding assay to determine antibody positive responses. Anti-RBD antibodies have lower cross-reactivity and background binding by ELISA compared to anti-spike antibodies. This results in a higher overall sensitivity and specificity for the assay. While the full-length spike protein has a larger number of epitopes that antibodies can bind which may correspond to higher antibody titers, other groups indicated that most individuals tested positive or negative for both anti-RBD and anti-spike antibodies with few individuals testing positive for one but not the other^30,31^. Similarly, our validation cohort showed that anti-RBD and anti-spike antibody results were congruent.

Serological protection conferred by vaccination was significantly more robust compared to antibodies induced by natural viral infection. Both total anti-RBD IgG binding, as well as *in vitro* neutralizing activity were stronger in vaccinated subjects^32^. The immune system may respond more efficiently to a single protein during vaccination, allowing for a more focused immune response to fewer epitopes, compared to the array of viral proteins and epitopes present during natural infection. Moreover, vaccination elicited higher antibody titers in participants who were pre-immune to SARS-CoV-2 compared to those who were immunologically naïve. This robust and immediate recall of high affinity antibodies may be attributed to memory B cell mediated processes^33^, similar to immune responses shown in other infectious agents^34,35^, as well as in antibody binding to SARS-CoV-2^33,36^ and in neutralization assays performed using a pseudo-typed virus displaying SARS-CoV-2 spike protein^23^. These findings highlight the importance of vaccination, especially in light of reports of short-term protection and abundant cases of reinfections after antibodies elicited by natural infection^37,38^.

A single vaccination with an mRNA vaccine was sufficient to significantly increase the neutralization titer in humans previously infected with SARS-CoV-2^39^. Anti-RBD antibody binding and VN supported the conclusion that all 8 pre-immune participants examined in this study had a significant increase in antibody levels prior to the administration of the second vaccination with no further significant change in antibody levels after the second vaccination. Not only did the first vaccination yield a more significant rise in antibody levels in pre-immune participants compared to immunologically naïve participants, pre-immune individuals had a more rapid rise following the first vaccination, with some individuals experiencing a significant increase in antibody titer in just 4 days after the first vaccination. Such a rapid increase in IgG levels in serum so soon after vaccination most likely indicates a memory cell-driven response that was recalled from a prior natural infection^40–42^.

Immunologically naïve participants could be categorized into two groups: individuals that seroconverted before the second vaccination, and those who did not seroconvert until after the second vaccination. This is likely due to the timing of the blood collection. In general, serum samples collected earlier after the first vaccination (average = 12 days, range = 9-18 days) had no change in antibody levels, while participants with later blood collections (average = 19 days, range = 12-26 days) tended to have an increase in antibody titers. There was a significant positive correlation between the number of days after the first vaccination that the serum was taken and the antibody level (Fig. S1A-B), while the same trend was less apparent in infected participants (Fig. S1C-D). Other research groups have shown similar trends because later serum collections allowed additional time for the immune system to mount a *de novo* response to the vaccine antigens^43^. However, there was no correlation between the number of days after the second vaccination and the titer level amongst immunologically naïve (Fig. S1E-F) or pre-immune participants (Fig. S1G-H).

Even though there was no significant change in antibody levels in pre-immune participants in response to the second vaccination, delaying or even skipping the second vaccination may have unknown, long-term effects that have not yet been explored. The longevity of antibodies or the quantity and quality of the memory B cells could be impaired compared to participants who received a full vaccine regimen, and there could be additional long-term advantages for fully vaccinated participants even when short-term serological benefits are not immediately apparent.

Overall, this study validated a robust ELISA assay for detecting SARS-CoV-2 anti-RBD IgG antibody binding with high sensitivity and specificity in human sera. Using a set of serum samples collected from 20 immunologically naïve and 20 pre-immune age-matched and vaccine-matched participants, it was demonstrated that vaccination with SARS-CoV-2 mRNA vaccines elicits higher serological binding and neutralizing antibody levels than non-vaccinated individuals who were naturally infected. Furthermore, a single vaccination was sufficient to boost pre-immune participants and the second vaccination did not further increase their antibody levels. In immunologically naïve participants, ~50% of participants had no significant rise in antibody titers until after the administration of the second vaccination, while the other half whose titers started to rise prior to the second vaccination were significantly boosted further by receiving the second vaccination. Future studies will assess the longevity and magnitude of anti-RBD and neutralizing antibody titers after vaccination between immunologically naïve and pre-immune individuals.

## Acknowledgments

The authors would like to thank the SPARTA collection and processing teams in Athens and Augusta, GA, as well as Debbie Bratt for program coordination. The authors also thank Katie Mailloux, Jasmine Burris, Omar Hamwy, Hua Shi, Naveen Gokanapudi, Lillian Buescher, Patrick Fagan, Brittany Baker, Charlotte Bolle, Courtney Briggs, Tejal Hill, Jordan Byrne, and Lauren Howland for technical assistance. We acknowledge the staff at the Animal Health Research Center at the University of Georgia for the upkeep and maintenance of the BSL-3 facilities. The recombinant proteins were produced by Jeffrey Ecker, Spencer Pierce, Ethan Cooper, and the team in the Center for Vaccines and Immunology protein production core. We would also like to thank all participants enrolled in the study, as well as Dr. Brad Phillips, Kimberly Schmitz, and the entire staff at the University of Georgia Clinical and Translational Research Unit (CTRU) for assistance in collecting samples in the SPARTA program. The CTRU was supported by the National Center for Advancing Translational Sciences of the National Institutes of Health under Award Number UL1TR002378.

This study was funded, in part, by the University of Georgia (US) (UGA-001), and by the National Institute of Allergy and Infectious Diseases (NIAID), a component of the U.S. National Institutes of Health (NIH), Department of Health and Human Services, under contract 75N93019C00052. In addition, TMR is supported by the Georgia Research Alliance as an eminent scholar (GRA-001). The content is solely the responsibility of the authors and does not necessarily represent the official views of the NIH.

## Supplemental Information

**Table S1. Demographic and serological information for the SPARTA participants in this study.** Dates for pre-, mid-, and post-vaccination timepoints are also noted. Antibody concentration refers to anti-RBD IgG antibodies, VN titer represents the reciprocal of the highest dilution point where cytopathic effects were not yet visible. (A) Participants from the immunologically naïve group who have no confirmed SARS-CoV-2 infection prior to vaccination. A single participant, P-032 had serum drawn between the two vaccinations. (B) Participants from the pre-immune group who have had confirmed SARS-CoV-2 infection prior to vaccination. 8 participants had serum drawn between the two vaccinations. (C) An additional 12 immunologically naïve participants who had serum collected between the two vaccinations.

**Table S2. Demographic and serological information for the validation cohort.** Antibody concentration refers to anti-RBD IgG antibodies, VN titer represents the reciprocal of the highest dilution point where cytopathic effects were not yet visible.

**Fig. S1: Antibody response based on number of days after the reception of the first and second vaccinations.** In immunologically naïve participants, (A) neutralizing and (B) anti-RBD IgG antibody levels showed a positive correlation with the number of days after the first vaccination the serum was collected (*r*=0.7781, ** *p*=0.0017, and *r*=0.6861, ** *p*=0.0096 respectively). In infected participants, (C) neutralizing antibody levels showed no correlation, with the number of days after the first vaccination the serum was collected (*r*=0.5319, *p*=0.1749), but (D) binding antibody levels did show a slight positive correlation (*r*=0.7085, * *p*=0.0492). In immunologically naïve participants, (E) neutralizing and (F) binding antibody levels showed no correlation with the number of days after the second vaccination the serum was collected (*r*=-0.3313, *p*=0.064, and *r*=-0.2190, *p*=0.2284 respectively). Similarly, in infected participants, neither (G) neutralizing nor (H) binding antibody levels showed any correlation with the number of days after the second vaccination the serum was collected (*r*=0.2068, *p*=0.3817, and *r*=-0.2458, *p*=0.2963 respectively). In order to represent the values on a logarithmic scale, lack of neutralization was reported as 1.

