## Supplementary figures and images for "Functional characterization of SARS-CoV-2 vaccine elicited antibodies in immunologically naïve and pre-immune humans"

### Fig. S1

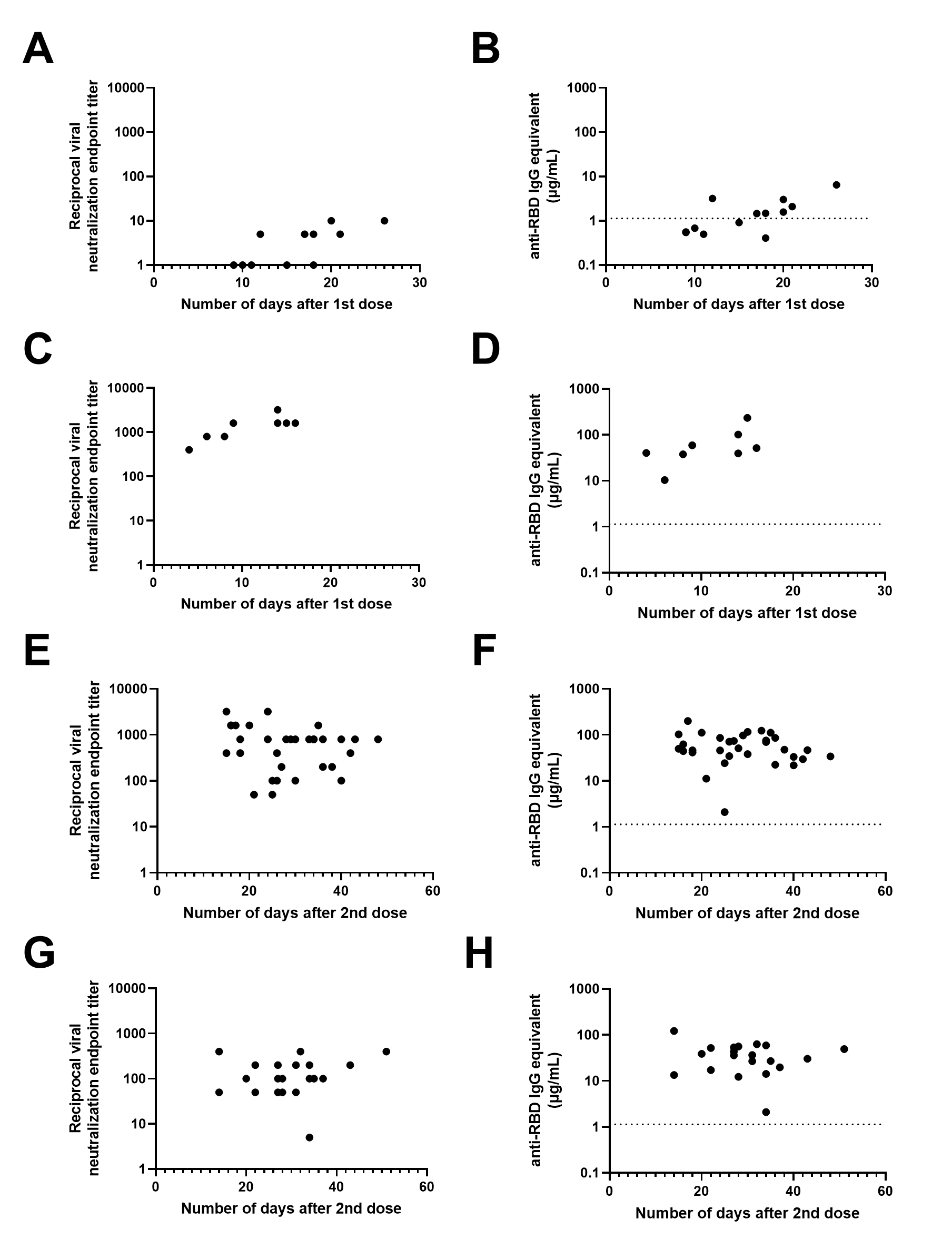
